# Neuroimmune cortical organoids overexpressing C4A exhibit multiple schizophrenia endophenotypes

**DOI:** 10.1101/2023.01.20.524955

**Authors:** Morgan M. Stanton, Sara Modan, Patrick M. Taylor, Harsh N. Hariani, Jordan Sorokin, Brian G. Rash, Sneha B. Rao, Alejandro López-Tobón, Luigi Enriquez, Brenda Dang, Dorah Owango, Shannon O’Neill, Carlos Castrillo, Justin Nicola, Kathy Ye, Robert M. Blattner, Federico Gonzalez, Dexter Antonio, Pavan Ramkumar, Andy Lash, Douglas Flanzer, Sophia Bardehle, Stefka Gyoneva, Kevan Shah, Saul Kato, Gaia Skibinski

**Author notes:** Primary Correspondence.

## Abstract

Elevated expression of the complement component 4A (C4A) protein has been linked to an increased risk of schizophrenia (SCZ). However, there are few human models available to study the mechanisms by which C4A contributes to the development of SCZ. In this study, we established a C4A overexpressing neuroimmune cortical organoid (NICO) model, which includes mature neuronal cells, astrocytes, and functional microglia. The C4A NICO model recapitulated several neuroimmune endophenotypes observed in SCZ patients, including modulation of inflammatory genes and increased cytokine secretion. C4A expression also increased microglia-mediated synaptic uptake in the NICO model, supporting the hypothesis that synapse and brain volume loss in SCZ patients may be due to excessive microglial pruning. Our results highlight the role of C4A in the immunogenetic risk factors for SCZ and provide a human model for phenotypic discovery and validation of immunomodulating therapies.

## Introduction

Schizophrenia (SCZ) is a psychiatric disorder characterized by psychotic and cognitive symptoms. While pathogenic mechanisms of SCZ remain unknown, specific disease pathology includes loss of gray matter, dendritic spines, and synapses, and increased cytokine expression^1–3^. Current treatments for SCZ address psychotic symptoms, but no existing therapies address the underlying disease pathology. Thus, a greater understanding of SCZ pathology in human models would provide valuable insight for development of disease modifying therapeutics.

The largest genetic risk factors for SCZ are associated with the immune-linked major histocompatibility complex (MHC) and complement component 4 (C4) genes consisting of C4A and C4B^4,5^. Increased copy number variants of C4A correlate with increased SCZ risk^5^, with elevated levels of C4A found in post-mortem brain samples^5^ and cerebrospinal fluid (CSF) of patients with SCZ^6^. Increased complement C3 deposition on synaptic structures, initiated by C4 in the complement cascade, play a pivotal role in microglia-mediated elimination of synapses^7,8^. Mouse models overexpressing C4A display a loss of synapses^9^ and increased phagocytosis of synapses by microglia^10,11^. Furthermore, copy numbers of C4A of microglia cultured in 2D correlated with an increase of phagocytosis of synapses^12^. These studies add to the growing hypothesis that elevated C4A in SCZ patients cause excessive synaptic pruning by microglia leading to reduced synaptic density and cognitive deficits.

The current lack of human *in vitro* SCZ models hinders mechanistic understanding of the disease and limits drug screening capabilities. Human iPSC-derived brain organoids offer the potential to understand disease etiology and discover novel therapeutics in a more biologically realistic system than monolayer cultures. Organoids retain the scalability of *in vitro* models for high throughput therapeutic screens and present a richer set of high-dimensional structural and functional phenotypic data modalities^13^. Brain organoids have a rich diversity of cell types that better replicate the *in vivo* CNS. Unlike 2D neuronal cultures, the complex tissue-like assembly of neurons and glia in 3D enables the capture of cell specific and cell-cell interactions during disease pathogenesis, including microglia-induced synaptic pruning.

Incorporation of microglia into organoids offers a novel axis to understand disease mechanisms associated with neuroimmune biology. Here, we present the first lentiviral (LV) overexpression C4A neuroimmune cortical organoid (NICO) platform for SCZ therapeutic discovery. NICOs contain robust populations of functional microglia, astrocytes, and neurons. Over-expression of C4A in the NICO model induces non-cell autonomous phenotypes including altered expression of neuroinflammatory genes, increased cytokine expression, and increased microglia phagocytosis of synapses. The C4A expressing NICOs capture multi-dimensional SCZ disease biology that have not previously been demonstrated in other *in vitro* models, opening the door for scalable human biology-led approaches to therapeutic discovery.

## Results

### Microglia integrated within NICOs display physiologically relevant features

We developed a protocol to integrate functional microglia into forebrain organoids to generate NICOs (Fig. 1). Cortical organoids (COs) were differentiated from iPSCs using a single-SMAD organoid protocol, confirming cortical fate with EMX1, CTIP2, and TBR1 neuronal staining (Supplemental Fig. 1). iPSC-derived microglia were added to day 65 or older COs which reliably include astrocytes and mature neurons (Fig 1a. & Supplemental Fig. 1). We found that microglia uniformly integrate into cortical organoids after 21 days of incubation (Fig. 1a & b, Supplemental Fig. 2 & Supplemental Video 1). We observed microglia morphology between 1 and 4 weeks post microglia addition to COs. After a four week incubation period, microglia showed greater ramification compared to microglia incubated for only 1 week, which had more ameoboid-like bodies (Fig. 1c). Extended culture of microglia-neuron 2D co-culture systems beyond 2-3 weeks is a challenge^14^. However, the NICO model was able to support functional microglia 60 days post integration into a CO (Supplemental Fig. 3), emphasizing the capabilities of the NICO for replicating *in vivo* neuroimmune biology. To quantify microglia morphology, NICOs from each incubation week post microglia addition were harvested, sectioned, and fluorescently labeled with the microglia marker IBA1. We employed machine learning-based segmentation analysis to create segmentation masks and highlight IBA1 positive areas within the NICO tissue (Fig. 1d). After applying IBA1 segmentation masks, we extracted geometric morphology statistics (Fig. 1e & f) for the microglia cell bodies. From 1 week to 4 weeks incubation, microglia exhibited an overall decrease in circularity and increase in perimeter, indicating increased ramification and fewer cells in an amoeboid state. The morphological changes observed with longer incubation periods of microglia in NICOs recapitulate changes observed with *in vivo* microglia as they migrate into the brain and become more ramified^15^ and suggest an appropriate period for assessing NICO immunological functionality.

**Figure 1.**
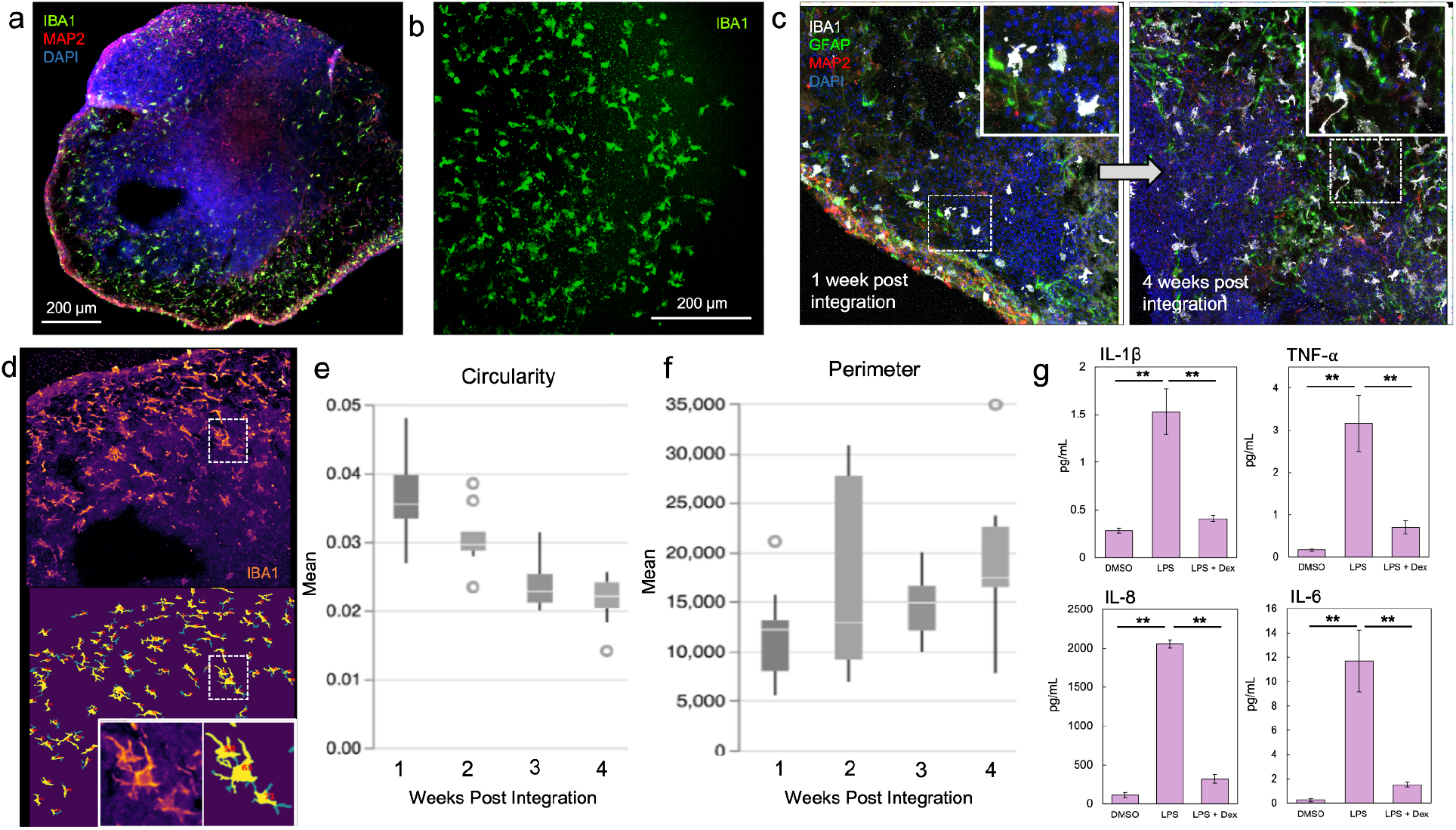
Microglia integrated within NICOs display physiologically relevant features. (**a & b**) Microglia were integrated into COs for 21 days to form NICOs. Immunostaining with microglia marker IBA1 demonstrates distribution throughout the CO tissue. (**b**) 3D projection of microglia (IBA1) throughout a whole cleared NICO. **(c**) Representative images of microglia (IBA1) integrated into NICOs after 1 or 4 weeks showing that prolonged incubation of microglia increases microglia ramification. White dashed boxes indicate microglia enlarged images in insets. (**d**) Examples of microglia masks generated from microglia segmentation analysis. (**e & f**) Quantification of microglia morphology after 1-4 weeks of integration into COs shows a decrease in microglia (e) circularity and an increase in (f) perimeter (*n* = 4-5 separate NICOs per parameter per time point, box plot = upper and lower quartiles, with median center line, error bars = max and min of data set, individual points represent data outliers.) (**g**) NICOs incubated with 100 ng/mL LPS show an upregulation of secreted proinflammatory cytokines, compared to NICOs treated with 100 ng/mL LPS and 10 µM dexamethasone (Dex) or 0.1% DMSO (*n* = 6-8 separate NICOs per parameter, bar = sample mean, error bars = SEM, statistical significance determined by one-way ANOVA with Tukey post-hoc tests, \*\**p* < 0.01; corrected for multiple comparisons).

To examine the general immune response of the microglia in NICOs, lipopolysaccharide (LPS) challenges were performed on day 92 NICOs, three weeks post microglia integration. NICOs were incubated overnight in 100 ng/mL LPS, 100 ng/mL LPS with 10 µM dexamethasone (Dex), or 0.1% DMSO and the media was collected for cytokine analysis (Fig. 1g). LPS elicited a strong microglial cytokine response with significant increases in secreted cytokines IL-1β, IL-8, IL-6, and TNF-α compared to the control. Addition of Dex, an anti-inflammatory corticosteroid, with LPS to the NICOs inhibited cytokine release (Fig. 1g). The ability to efficiently modulate and quantify the inflammatory state of the NICO system provides a powerful tool for neuroimmune disease phenotyping and therapeutic discovery assays.

### SCZ associated C4A expression in human NICOs

To selectively drive C4A expression in neurons, C4A was expressed under a synapsin promoter and delivered to day 63 COs using lentiviral (LV) delivery (Fig. 2a & b). To enable live cell visualization of expression, C4A was tagged to green lantern GFP (C4A-GFP) via a self-cleaving T2A tag. As a control, COs were also transduced with synapsin-driven GFP. Seven days post-transduction, we observed expression of the LV reporters in GFP and C4A-GFP COs via live fluorescence imaging (Fig. 2c). To validate expression of C4A, day 92 COs were harvested and quantified for C4A expression levels via RT-qPCR. COs overexpressing C4A-GFP showed significantly higher C4A levels compared to COs transduced with GFP or controls with no lentivirus added (No LV) (Fig. 2d). Additionally, secreted media was harvested and quantified for levels of C4. NICOs overexpressing C4A-GFP showed higher secreted C4 levels compared to COs transduced with GFP or with no LV (Supplemental Fig. 4). Following confirmation of C4A expression and secretion, microglia were incubated for 3 weeks with transduced COs to generate C4A-GFP and GFP NICOs. C4A-GFP NICOs exhibited reproducible brightfield morphology that showed no organoid body morphological differences when compared to NICOs overexpressing GFP or with No LV (Fig. 2e). We quantified microglia integration rates via flow cytometry analysis of day 91 C4A-GFP NICOs and found that C4A overexpression did not inhibit microglia integration into COs (Fig. 2f). GFP and C4A expression were also durable after microglia integration (Fig. 2g & h respectively). 91 day old NICOs expressing C4A maintained significantly higher C4A transcripts compared to GFP only or no LV NICOs (Fig. 2h) and levels were similar to that seen in COs prior to microglia addition.

**Figure 2.**
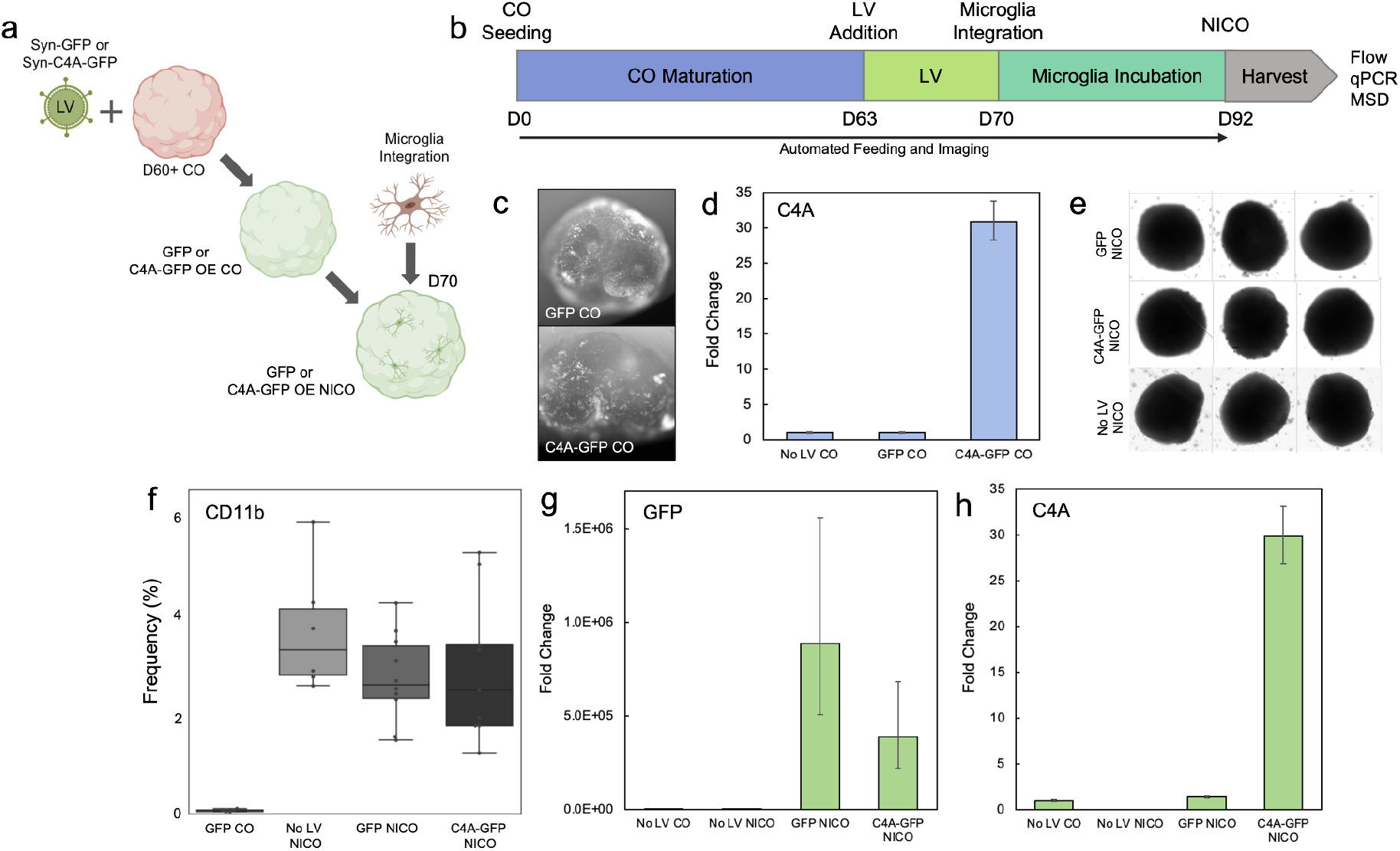
SCZ-associated C4A expression in human NICOs. (**a & b**) Summary and timeline of generation of C4A NICOs. Day 63 cortical organoids (COs) were transduced with lentivirus (LV) expressing C4A-GFP or GFP driven by a synapsin promoter. 7 days post-transduction microglia were integrated into COs to form C4A-GFP or GFP NICOs. (**c**) Live imaging shows GFP fluorescence in C4A-GFP and GFP day 70 COs, 1 week post LV delivery. (**d**) RT-qPCR quantification of C4A of day 92 in C4A-GFP or GFP, or No LV COs. (**e**) Representative bright field images of day 92 NICOs overexpressing C4A-GFP, GFP, or with No LV. (**f**) Frequency of microglia marker, CD11b in day 92 NICOs overexpressing C4A-GFP, GFP, or with No LV. A GFP CO is also included as a negative control (*n* = 6-8 separate organoids per parameter, box plot = upper and lower quartiles, with median center line, error bars = max and min of data set, statistical significance determined by one-way ANOVA with Tukey post-hoc tests, no significant difference in data between NICO groups). (**g & h**) RT-qPCR quantification of (g) GFP and (h) C4A of day 92 No LV COs and No LV, GFP, and C4A-GFP NICOs (For all qPCR, *n* = 4-5 pooled COs or NICOs per parameter run in triplicate, qPCR relative to GAPDH, fold change relative to No LV CO, bar = sample mean of technical replicates, error bars = s.d. of technical replicates).

### C4A expression in NICOs modulates proinflammatory markers that are associated with SCZ

Neuroinflammation is a common pathophysiology of SCZ, with patients showing elevated cytokine and chemokine responses and activated microglia markers^16,17^. To test if elevated C4A levels could directly induce inflammatory responses in human-derived organoids, C4A-GFP and GFP NICOs and their media were harvested for expression of inflammatory markers and secreted cytokines. A selection of 12 microglia genes previously linked with disease associated microglia (DAM), Alzheimer’s, or SCZ were analyzed via RT-qPCR (Supplemental Table 1)^18–21^. NICO samples were pooled and lysed for cDNA. Four genes of the 12 tested were significantly up- (B2M) or down-regulated (CST7, ITGAX, and LIPA) relative to the GFP control group (Fig. 3a). At least one of these genes, ITGAX, was previously shown to be down-regulated in SCZ patients^19,22^. Previous comparisons of inflammatory genes associated with Alzheimer’s and SCZ have shown opposite effects, where genes up-regulated in Alzheimer’s were down-regulated in SCZ and vice versa^19^, suggesting specific neuroinflammation pathways for each pathology. We found opposing effects of the phagocytic and lipid metabolism gene, CST7, which demonstrated up-regulation in an Alzheimer’s mouse model^18^, but down-regulation in our system.

**Figure 3.**
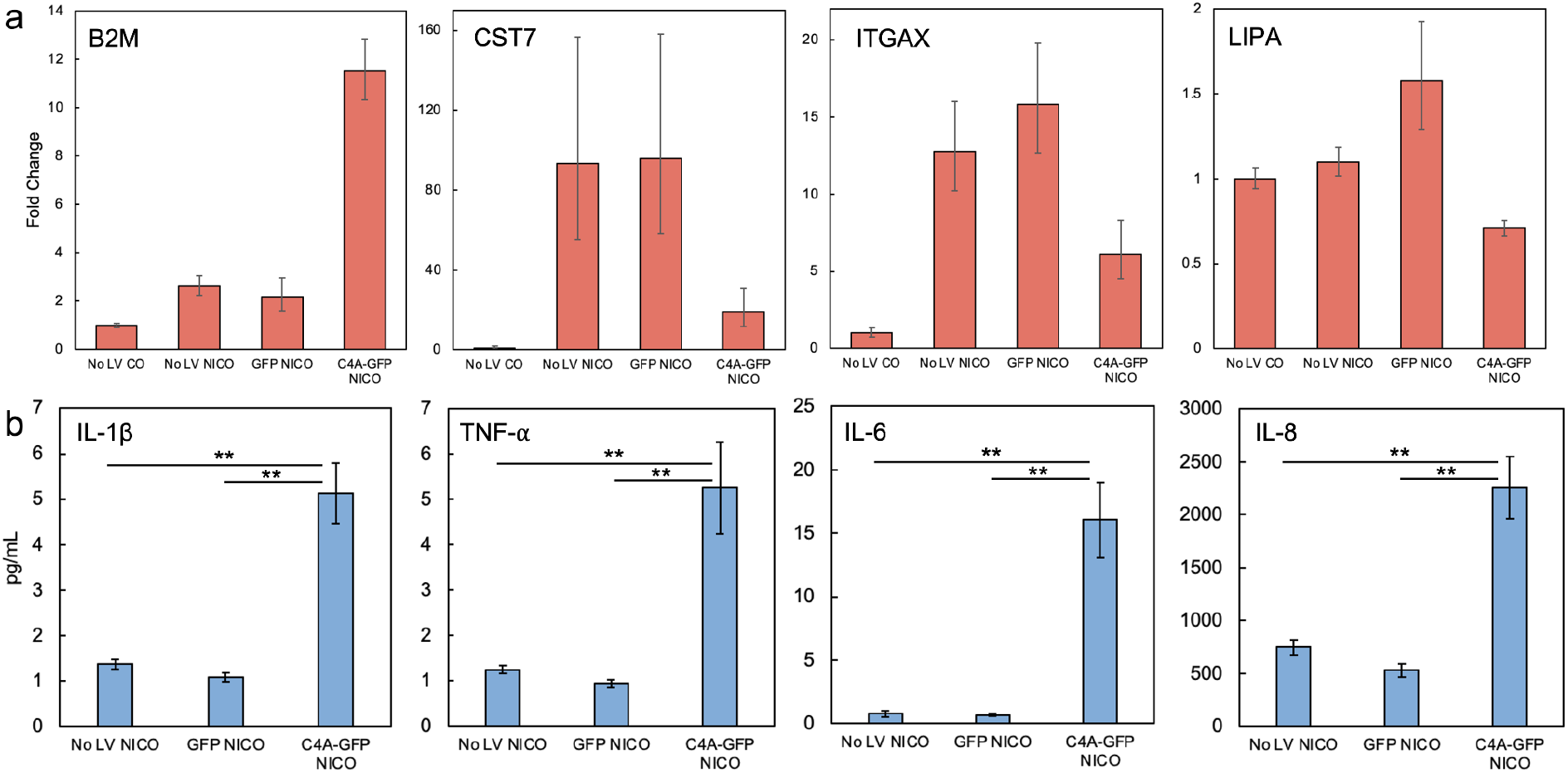
C4A expression in NICOs modulates proinflammatory markers. (**a**) RT-qPCR quantification of up-regulated B2M and downregulated CST7, ITGAX, and LIPA in day 92 No LV COs and No LV, GFP, and C4A-GFP NICOs (For all qPCR, *n* = 4-5 pooled COs or NICOs per parameter run in triplicate, qPCR relative to GAPDH, fold change relative to No LV CO, bar = sample mean of technical replicates, error bars = s.d. of technical replicates). (**b**) Levels of secreted IL-1β, TNF-α, IL-6, and IL-8 cytokines in media from NICOs expressing C4A-GFP or GFP, or No LV, (*n* = 8-10 independent samples per condition, bar = sample mean, error bar = SEM, statistical significance determined by one-way ANOVA with Tukey post-hoc tests, \*\**p* < 0.01; corrected for multiple comparisons).

To assess secreted cytokines, media was collected from C4A-GFP and GFP NICOs and analyzed for levels IL-1β, TNF-α, IL-6, and IL-8 cytokines. Significant increases in all four secreted cytokines were found in C4A-GFP NICOs compared to GFP and no LV NICOs (Fig. 3b). Minimal cytokine release was observed in GFP and GFP-C4A COs (Supplemental Fig. 5), indicating microglia were the primary driver for C4A-induced cytokine secretion. The cytokine release in the C4A NICOs represents a heightened neuro-inflammatory response in the microglia similar to what is seen in the CSF ^23,24^ and plasma^25^ of SCZ patients. Our cytokine and gene expression results of the C4A-GFP NICO model highlight the ability of the model to recapitulate SCZ specific patient neuroinflammatory pathology.

### C4A expression in human NICOs promotes synaptic phagocytosis

C4 has been implicated with other complement proteins to promote synaptic pruning^10,11^. C4A overexpression observed in C4A patients is proposed to contribute to the decrease in synapse numbers observed in SCZ patients^26,27^. A flow cytometry-based assay was developed to detect intracellular synaptic proteins in microglia in C4A-GFP NICOs. Similar methods are previously described by Yilmaz *et al*.^*11*^ and Makinde *et al*.^*28*^ to quantify microglia phagocytosis in mice. Here, microglia were harvested from NICOs, stained for CD11b and the presynaptic marker, synaptic vesicle glycoprotein 2A (SV2A) and analyzed via flow cytometry for detection of intracellular synaptic proteins (Fig. 4a & Supplemental Fig. 6 a & b). Microglia cultured in C4A-GFP NICOs had significantly more synaptic content than microglia from GFP NICOs as shown by mean fluorescent intensity (MFI) (Fig. 4b), demonstrating an increase in microglial phagocytic activity in the presence of C4A. Furthermore, the percentage of microglia co-labeled with SV2A did not change between GFP and C4A-GFP NICOs (Supplemental Fig. 6c) indicating microglia had more synaptic material per cell in the C4A model than controls. To strengthen this finding, a live imaging-based assay was developed to capture the dynamic process of microglia phagocytosis within COs. RFP labeled microglia were integrated into the C4A-GFP and GFP COs (Fig. 4c). One to two weeks after microglia integration, a live imaging time series was used to monitor microglia phagocytosis in GFP NICOs (Fig. 4d & Supplemental Video 2) and C4A-GFP NICOs (Fig. 4e & Supplemental Video 3). Microglia cultured in C4A-GFP NICOs had higher GFP payload compared to microglia in GFP only organoids. These results support the hypothesis that elevated C4A leads to microglia-mediated elimination of synapses, which would in turn drive over-pruning of synapses in SCZ patients.

**Figure 4.**
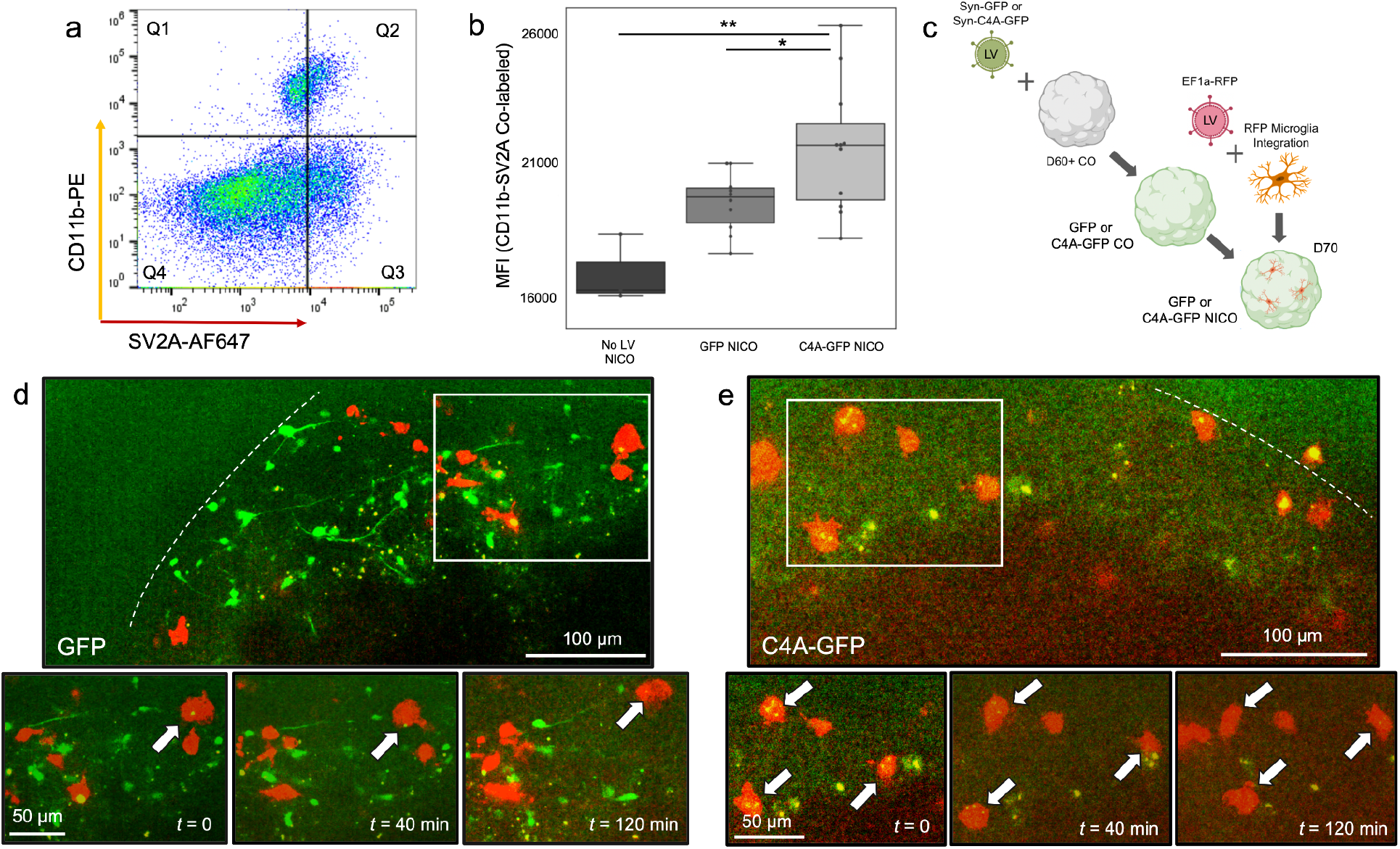
C4A expression promotes synaptic phagocytosis. (**a & b**) C4A-GFP and GFP NICOs were harvested and quantified for synapse engulfment by microglia using flow cytometry co-staining of microglia (CD11b) and presynaptic protein SV2A. The gating strategy is shown in (a). (**b**) The CD11b-SV2A mean fluorescence intensity (MFI) in Q2 of No LV, GFP, and C4A-GFP NICOs (*n* = 4-12 separate organoids per parameter, box plot = upper and lower quartiles, with median center line, error bars = max and min of data set, statistical significance determined by one-way ANOVA with Tukey post-hoc tests, \**p* < 0.05, \*\**p* < 0.01; corrected for multiple comparisons,). (**c**) Summary for labeling microglia with RFP fluorescent LV to monitor live phagocytosis in GFP and C4A-GFP NICOs. (**d & e**) Live imaging time series of RFP labeled microglia ingesting GFP labeled neurons in (d) GFP and (e) C4A-GFP NICOs. White dashed line outlines the NICO edge. White box indicates microglia tracked during time series. White arrows indicate co-localization of microglia (RFP) with GFP labeled phagocytic cargo.

## Discussion

Previous studies have integrated microglia into 3D neuronal cultures using various strategies^29^, and in contrast to 2D and mouse models, microglia cultured in 3D remain in a more stable homeostatic state^30^. However, achieving long term organoid-microglia co-cultures at scale has been a challenge for the field. COs generated via neuroectoderm induction using dual SMAD inhibition do not contain endogenous microglia; therefore we developed a NICO protocol that enables reproducible integration of human iPSC-derived microglia into COs. Using our automated CO culture and imaging platform^13^, we established a highly reproducible human NICO model with functional microglia (Fig. 1 & Supplemental Fig. 2). To demonstrate functionality of the NICO model we cultured microglia over an extended period and recorded shifts in morphology towards a more ramified state, similar to the morphological changes observed when microglia migrate from the embryonic yolk sac into the brain during development. We also observed during LPS challenge, a common method for inducing microglial transition into an inflammatory state, that NICOs elicited increased cytokine expression (Fig. 1g). Using this model, we demonstrate that C4A overexpressing NICOs can recapitulate an array of SCZ-associated phenotypes along the neuroimmune axis, providing a platform for novel SCZ therapeutic discovery and validation.

Proinflammatory cytokine expression can be monitored as a hallmark disease phenotype as continually activated microglia become detrimental to neuronal health^31^. Chronically activated microglia and cytokine expression have been associated with SCZ, multiple sclerosis, Alzheimer’s and Huntington’s disease and linked to excessive synapse pruning, neuronal death, and astrogliosis^6,32^. Overexpression of C4A in human NICOs does not hinder microglia integration (Fig. 2f); however, microglia move into a chronic proinflammatory state while exposed to excessive C4A (Fig. 3b). The increase in cytokine levels induced by C4A overexpression matches increases in cytokine levels seen in patients with SCZ, and is important because patient levels of cytokine expression directly correlate with patient SCZ severity and duration of the disease. Higher levels of IL-6, IL-1β, and TNF-α are linked to longer hospital stays and more severe SCZ symptoms^1^, while increased levels of IL-1β correlate with decreased verbal fluency and a reduction in brain volume^33^. These findings suggest that microglia inflammatory state and cytokine levels play a direct role in SCZ and are an early indicator for patient psychosis^34^.

One major clinical phenotype associated with SCZ is loss of gray matter and brain volume that is observable in SCZ patients as early as their first episode^35^. This loss has been associated with excessive microglia-mediated synaptic pruning^36^. C4 is a primer step in the immune complement cascade to initiate synaptic pruning as C4 promotes C3 binding to synaptic targets and initiates their engulfment by microglia^5^. Using our NICO C4A overexpression model, we developed a phagocytic activity assay to quantify microglia uptake of the presynaptic marker SV2A (Fig. 4a & b). SV2A is pervasive throughout the brain, is localized to GABAergic and glutamatergic presynaptic nerve terminals, and plays a role for neurotransmitter exocytosis^37^. Changes in SV2A also have functional consequences, including abnormal inhibitory neurotransmission^38^. We find that microglia in C4A-GFP NICOs have significantly increased levels of SV2A, consistent with other reports of C4A overexpression systems showing increased uptake of synaptic material in mice^9–11^ and 2D microglia^12^. Recently, decreases in SV2A volume in the frontal cortex of SCZ patients have been observed when compared to healthy volunteers and may represent the basis for decreased gray matter and brain volume seen in SCZ patients. However, there were no correlations between symptom severity and decreased volume, suggesting that SV2A changes occur in earlier neurodevelopment^39^. Our work suggests that SV2A may be an important early biomarker for the presence and progression of SCZ.

Current treatments for SCZ rely on antipsychotics for modulation of dopaminergic circuits, but antipsychotic treatment does not assist with cognitive symptoms of the disease, does not address underlying disease mechanisms, nor does it prevent further neural impairment^40^. The C4A NICO is the first of its kind human *in vitro* SCZ model capable of replicating human biology not observed in 2D or animal systems. The C4A NICO model enables the study of multiple SCZ phenotypes concurrently and offers a targeted approach for testing pharmacological therapeutics in human tissue. In our reproducible and scalable human C4A NICO model, treatments can be assessed for their ability to rescue synapse elimination by microglia, cytokine expression and changes in proinflammatory gene states. Our results support the hypothesis that the neuroimmune axis plays a critical role in the development of SCZ and provides a platform for SCZ patient-specific drug discovery.

## Materials and Methods

### Cell lines used for generating COs and microglia

A control donor iPSC clone obtained from CIRM (CW20109) that had no history of neuropsychiatric disease and no known schizophrenia-associated gene variants was used for organoid and microglia differentiation.

### Culture of cortical organoids (COs)

COs were grown in individual wells using a novel dorsal forebrain protocol derived from Velasco *et al*.^41^ to improve cortical fate, reliability, and embryoid body survival in round bottom plates. Briefly, iPSCs were thawed concurrently and passaged twice before seeding. Prior to seeding, iPSCs were dissociated when 50–80% confluent and plated in U-bottom 96-well plates at 9×10^3^ cells per well with ROCK inhibitor. Organoids were cultured in neural induction media containing inhibitory molecules added to direct cortical fate via single-SMAD, Wnt signaling inhibition, as well as Shh inhibition. Organoids were fed on automated Biotek feeders and washers. On the fifth week of culture, organoids were switched to a DMEM-based maturation media with the addition of Matrigel Growth Factor Reduced (GFR) Basement Membrane Matrix. On the eighth week of culture, COs were fed maturation medium containing BDNF and GDNF with GFR Matrigel. Matrigel was omitted during the ninth week of culture. COs were maintained with automated feeding and imaging until harvest for LV transduction and microglia integration.

### Differentiation of microglia from iPSCs

Microglia were differentiated from the same iPSCs used to generate COs. To induce mesoderm development, iPSCs were treated with BMP4, CHIR99021 and bFGF for approximately two to three days. To induce hematopoietic progenitors, cells were treated with TPO, VEGF, SCF, and IL-6 for an additional five to six days. To generate microglia, cortical hematopoietic progenitors were plated on matrigel-coated flasks with IL-34, M-CSF, and TGF-1 in serum-free conditions for three weeks. Cells were collected and stored at -80 °C until use. To confirm microglia differentiation, cells underwent in-house quality control (QC) verification. Briefly, a sample of the microglia population was thawed and accessed for (1) viability greater than 80% post thaw, (2) positive staining of CD11b (PE-rat, Miltenyi, 130-113-235, 1:50), CD45 (FITC-mouse, BD, 555482, 1:100), TMEM119 (mouse primary, Biolegend, 853302, 1:250, donkey anti-mouse 488 Life Technologies, A21202, 1:1000), and CX3CR1 (PE-rat, Thermo Fisher Scientific, 12-6099-42, 1:20) via flow cytometry, (3) positive staining for IBA1 (rabbit primary, Wako, 019-1974, 1:2000, donkey anti-rabbit 647 Life Technologies, A-31573, 1:1000) via 2D immunofluorescence imaging and (4) cytokine response to LPS challenges. Flow cytometry, immunofluorescence imaging, and cytokine measurements described in subsequent Methods paragraphs. Microglia were only used for experiments after QC verification of each production batch.

### Plasmid and lentivirus (LV) generation

To generate the C4A plasmids, human C4A transcript was synthesized based on the open reading frame from NM_001252204.2, which has previously been used to model C4A in mice^10^. DNA fragment synthesis and cloning were performed by GenScript. pLV-hSyn-glGFP were generated by removing the AgeI-RFP-PmeI insert of pLV-hSyn-RFP and ligating an AgeI-green lantern GFP -PacI-PmeI synthetic fragment. pLV-hSyn-C4A-T2A-glGFP were generated by removing the AgeI-RFP-PmeI insert of pLV-hSyn-RFP and ligating in a single reaction an AgeI-AscI-C4A-SpeI and an SpeI-T2A-glGFP-PacI-PmeI synthetic fragments^42^. The final constructs, pLV_hSyn_glGFP and pLV_hSyn_C4A-T2A-glGFP, were sequenced to confirm integration of the transgenes C4A and glGFP. Lentiviral production of pLV_hSyn_glGFP (referred to as ‘GFP’ in text) and pLV_hSyn_C4A-T2A-glGFP (referred to as ‘C4A-GFP’ in text) viral particles were generated by Signagen.

### Lentiviral transduction of organoids

For virus addition to day 63 organoids, all media was removed from organoid wells and replaced with 100 µL Maturation media containing Syn-C4A-GFP or Syn-GFP with 8 µg/mL polybrene (EMD Millipore, TR-1003-G). The final virus concentration was 10 MOI. Control groups without virus were seeded with media containing only 8 µg/mL polybrene. Organoids were incubated in virus-containing media overnight at 37 °C. After 24 hours, all virus-containing media was removed and 200 µL of fresh Maturation media was added to each well. For the following feeds, organoids had 65% media changes every day after virus addition with fresh Maturation media until organoids were harvested for analysis.

### Microglia integration for NICO formation

Twenty four hours before microglia integration, day 69 COs had 65% of their media removed from each well and replaced with MicroMaturation media (media recipe modified from Abud *et al*.^*43*^, maturation media supplemented with IL-34, TGFβ, M-CSF, and chemically defined lipid mix). COs were incubated in MicroMaturation media overnight at 37 °C. The following day, all media was removed from CO wells and replaced with 50 µL fresh MicroMaturation media containing 2×10^6^ microglia/mL per well. COs were incubated in Micro-Maturation media for 1 hour at 37°C followed by addition of 170 µL MicroMaturation media to each well. NICOs were incubated at 37 °C and 5% CO_2_ with 65% MicroMaturation media changes every other day until harvest.

### Immune functional microglia in NICOs

For testing immune functionality of microglia in NICOs, all media was removed from day 91 COs and NICOs. Media was replaced with fresh MicroMaturation media containing either 0.1% v/v DMSO, 100 ng/mL lipopolysaccharides (LPS, Sigma Aldrich, L6529), or 100 ng/mL LPS with 10 µM water soluble dexamethasone (Sigma Aldrich, D2915). COs and NICOs were incubated with compound solutions overnight at 37 °C and 5% CO_2_. The next day, media was collected, flash frozen on dry ice and stored at -80 °C until use.

### Reverse transcription (RT)-qPCR

Four to five organoids were physically lysed using a pipette and purified for RNA on the Promega Maxwell RSC48 in conjunction with the Maxwell RSC Simply RNA Tissue kit (Promega, AS1340). cDNA synthesis was performed using the SuperScript III First-Strand Synthesis System (Invitrogen, 18080051) in order to obtain a final cDNA concentration of 5 ng/µL. Manufacturer instructions were followed for both RNA purification and cDNA synthesis. RT-qPCR was executed in a semi-automated fashion and each condition was run as a technical triplicate. cDNA from GFP, C4A-GFP, or no LV NICOs were combined with QuantaBio Perfecta SYBR Green SuperMix Low ROX (QuantaBio 95056-02K). cDNA from COs with no LV were used as a negative control. GAPDH primers were used as an endogenous control, and exon spanning primers tested included GFP, C4A, B2M, CST7, ITGAX, CX3CR1, and LIPA (Supplemental Table 1). Primers and sample mixes were added to assay plates in triplicate. Plates were sealed, vortexed, and spun down before running on the Applied Biosystems QuantStudio 5 Real-Time PCR System. Analysis was completed using the QuantStudio Design & Analysis Software. The ΔΔCt method was used to represent fold change^44^.

### Cytokine quantification

Media samples were collected from individual wells of day 92 NICO and CO, flash frozen on dry ice, and stored at -80 °C. Upon use, samples were thawed on ice and used with the V-PLEX Human Proinflammatory Panel II (4-Plex) (Mesoscale Discoveries (MSD), K15053D-1) following manufacturer’s instructions. The assay plate was washed 3 times with 150 µL per well of phosphate buffered saline with 0.1% tween, pH 7.4 (PBST, BioWorld, 41620004), and dried. 50 µL of thawed media samples or standard MSD calibrators were added to each well. The plate was sealed and shaken at 750 rpm for 2 hours at room temperature. After incubation, the plate was decanted, washed 3 times with PBST, dried, followed by 25 µL of MSD antibody cocktail addition to each well. Antibody cocktail was prepared at a 1:50 dilution (MSD provided Sulfo-Tag detection antibodies: MSD Diluent 3). The plate was sealed and incubated at room temperature on a plate shaker at 750 rpm for 2 hours. After incubation, the plate was washed 3 times with PBST, dried, and 150 µL of 2X MSD Read Buffer T was added to each well. Read Buffer T was prepared by diluting Read Buffer T in equal parts deionized water. The plate was then immediately read on a MSD Quickplex SQ120 plate imager. Resulting data was analyzed using the MSD Workbench program and standard curve figures and calculated concentration data exported.

### Live brightfield and immunofluorescence imaging

Automated brightfield imaging of COs and NICOs was acquired using an in-house imaging system with a Flir Oryx 10GigE camera, Precise Automation PF-400 robotic arm, and images taken at 2x magnification. Live imaging of GFP or RFP immunofluorescence in COs and NICOs was obtained using IN Cell Analyzer 2200 (GE Healthcare) at 4x or 10x magnification. Videos of RFP microglia had images acquired every 10 minutes.

### Live immunofluorescence microglia labeling

For microglia fluorescence RFP labeling, microglia were thawed onto Matrigel-coated plates, transduced at 2 MOI with EF1α-TagRFP lentivirus (Cellecta pRSGERP-U6-sg-EF1-TagRFP-2A-Puro) in microglia culture media, collected after 48 hours, and integrated into organoids as indicated previously. NICOs were imaged 1-2 weeks after microglia integration.

### Fixed immunofluorescence sectioning, staining and imaging for NICO sections

COs and NICOs were transferred and rinsed with PBS and fixed in 4% paraformaldehyde (Electron Microscopy Sciences, 15710) for 4 hours at 4 °C. After fixation, organoids were rinsed 3 times with PBS, and incubated with 30% w/v sucrose solution overnight with gentle agitation at 4 °C. Samples were embedded into Optimal Cutting Temperature solution (Tissue Tek, 4583), flash frozen on dry ice, and stored at -80 °C until sectioning. Embedded blocks were sectioned into 25 μm slices using an Epredia NX-70 Cryostar Cryostat (Thermo Fisher). For staining, slides were rinsed with PBS, then incubated in 0.25% Triton X-100 with 4% Donkey serum (EMD Millipore, S30) for 1 hour, followed by incubation in primary antibodies overnight: rabbit IBA1 (1:400, Wako, 019-1974), mouse MAP2 (1:1000, Millipore-Sigma, MAB3418) and chicken GFAP (1:1000, Abcam, ab4674). After rinsing 5 times with PBS, slides were incubated in secondary antibodies, donkey anti-rabbit 647 (1:1000, Life Technologies, A-31573), donkey anti-mouse 555 (1:1000, Life Technologies, A31570), donkey anti-chicken 488 (1:1000, Biotium, 20166) and DAPI (1:1000, Biovision, B1098-5) for 2 hours at room temperature and rinsed again 5 times with PBS. Slides were then mounted and imaged using Zeiss Axio Scan Z1 with 20x / 0.8 objective and stitched together by the Zeiss Zen software.

### IHC segmentation analysis

NICO sections were imaged at 20x using an ImageXpress Confocal microscope (Molecular Devices). Microglia were segmented from a *z*-stack acquisition of NICO sections as follows. First, the maximum intensity projection across *z* = 4 planes was computed. Next, we log-transformed the IBA1 channel to enhance the intensity of microglia processes relative to cell bodies. To further enhance filaments, the log-transformed intensity image was blended with a Frangi vessel-transformed intensity image using an 80:20 ratio. Individual objects were segmented using a local Otsu threshold. Finally, we applied a graph-theoretic method to merge objects representing distal processes to objects representing their parent cell bodies and computed the perimeter and circularity metrics (*Circularity = 4*π *\* ((Area)/(Perimeter)*^*2*^)) as quantitative measures of microglia morphology. A circularity value of 1 designates a perfect circle, values below 1 indicate less circularity.

### Fixed immunofluorescence staining, clearing, and imaging for whole NICOs

NICOs were harvested from culture plates and fixed in 4% paraformaldehyde (Electron Microscopy Sciences, 15710) for 4 hours at 4 °C with gentle rocking. After fixation, NICOs were run through a 3D staining protocol that included: 5 washes with PTwH (PBS, 0.1% Tween-20, and 0.01% Heparin), overnight incubation in Perm Solution (PBS, 0.1% Tween-20, 0.1% Triton-x 100, 0.1% NP40, and 1 mg/mL Sodium deoxycholate) and followed by 5 washes with PTwH. The organoids were then placed in an overnight incubation with blocking solution (PBS, 0.1% Tween-20, 0.1% Glycine, 10 % DMSO, 5% donkey serum, and 5% BSA). Primary antibody (goat IBA1, Abcam, ab5076, 1:300) was added and the plate was placed on a tilt shaker (200 rpm for 72 hours, 4 °C), washed 10 times with PTwH for 10 minutes and then incubated with secondary antibody (donkey anti-goat AF647, Life Technologies, A21447, 1:1000) (and DAPI on a tilt shaker (200 rpm for 24 hours). At the end of secondary incubation, NICOs were washed (10 times for 5 minutes RT incubations between the first 5 washes and 30 minutes room temperature incubations between the last 5 washes, with PTwH) and moved to clearing or stored in PTwH and 0.05% sodium azide at 4 °C for up to 3 days before clearing.

NICOs were transferred to 96 well imaging plates (CellVis, P96-1.5P) and cleared by adding 80 µL Rapiclear 1.49 reagent (Sunjin Lab, RC149002) for 3 hours. NICOs were imaged on a ImageXpress Confocal High-Content Imaging System (Molecular Devices). Acquisitions were made automatically with a Objective 10x Ph1 Plan Fluor DL 0.30 (higher contrast) at channel-dependent optimized exposure times (1000 - 2000 ms), 77 z-stacks of 3 um were acquired per NICO.

### Flow cytometry

Organoids for flow cytometry had media removed and washed once with PBS. Organoids were dissociated into single cells using a 1:20 mixture of DNAse in PBS (Worthington Biochemical, LK003163) and TrypLE (Thermo Fisher, 12604013) respectively at 37 °C for 20 minutes followed by gentle manual dissociation with a pipette. An equal volume of 10% v/v fetal bovine serum (FBS, VWR, 45001-108) in PBS was added to the solution to quench the reaction. Cells were centrifuged at 300 rpm and washed 1 time with PBS, placed on ice, and labeled with Live-or-dye 750/777 stain (Biotium, 32008, 1:1000) and CD11b-PE rat antibody (Miltenyi, 130-113-235, 1:50) in cold PBS for 15 minutes. Cells were centrifuged and washed with PBS. Cells were fixed in 2% paraformaldehyde (Electron Microscopy Sciences, 15710) in cold PBS for 20 minutes at 4 °C followed by centrifugation at 1000 rpm, filtration through a 40 µM filter, and wash in PBS. Cells were centrifuged and resuspended in flow buffer for intracellular staining (buffer stock diluted in deionized water (Biolegend, 421002-BL) supplemented with 0.3% v/v BSA (Sigma-Aldrich, A9576), 1% v/v donkey serum (Sigma-Aldrich, S30-100ML) and 0.01% w/v sodium azide) and SV2A primary rabbit antibody (Abcam, ab254351, 1:1000). Cells were incubated in antibody solution overnight at 4 °C. Cells were centrifuged and washed 2 times in cold flow buffer and resuspended in donkey-rabbit AF647 secondary antibody (Life Technologies, A31573, 1:1000) and DAPI (Biovision, B1098-5, 1:10,000) in flow buffer. Cells were incubated for 30 minutes at room temperature, washed 2 times with flow buffer, resuspended in PBS, and stored at 4 °C until flow analysis. Flow cytometry was run on a ThermoFisher Attune NxT with auto sampler and Attune NxT v3.1.2 software. FlowJo v10.6.1 Software was used for data analysis.

## Statistical analysis

For all flow cytometry, cytokine, and ELISA data, statistical significance of differences between groups was determined using one-way ANOVA with Tukey post-hoc tests. Tests were deemed to be statistically significant at \**p* < 0.05 or \*\**p* < 0.01 levels, adjusted for multiple comparisons using Family-wise error rate.

## Author Contributions

MM Stanton, G Skibinski, K Shah, S Bardehle, S Gyoneva, and S Kato conceived and designed experiments. MM Stanton, K Shah, S Modan, PM Taylor, HN Hariani, BG Rash, SB Rao, A López-Tobón, L Enriquez and RM Blattner performed laboratory experiments. F Gonzalez advised on lentivirus design. D Owango, B Dang, K Ye, S O’Neill, C Castrillo, and J Nicola ran laboratory organoid production pipeline. J Sorokin, D Antonio, and P Ramkumar built machine learning models. J Sorokin, D Antonio, P Ramkumar, PM Taylor, S Modan, MM Stanton, and K Shah ran analysis. A Lash and D Flanzer built Herophilus laboratory software pipeline. C Castrillo, J Nicola, and D Flanzer built Herophilus automation imaging and feeding infrastructure. MM Stanton and G Skibinski wrote the manuscript with input from K Shah, S Kato, S Bardehle, and S Gyoneva. All authors discussed the results and reviewed the manuscript.

