## Supplemental Information for "Neuroimmune cortical organoids overexpressing C4A exhibit multiple schizophrenia endophenotypes"

### Supplemental Videos

**Supplemental Video 1.** Fly-through video of IBA labeled microglia in a whole cleared NICO.

**Supplemental Video 2.** Syn-GFP NICO with RFP labeled microglia. Images acquired every 10 minutes. Video at 2400x speed.

**Supplemental Video 3.** Syn-GFP-C4A NICO with RFP labeled microglia. Images acquired every 10 minutes. Video at 2400x speed.

### Supplemental Methods

### C4 ELISA

Media samples were collected from individual wells of C4A overexpressing COs culture plates on organoid day 92, flash frozen on dry ice, and stored at -80 °C. Upon use, samples were thawed on ice and used with the Human complement C4 Elisa Kit (Abcam, ab108825) following manufacturer's instructions. Briefly, the media was diluted 1:1 with PBS. 50 µL of each sample well was added to a microplate well with Complement C4 standards and incubated for 2 hours. The microplate was washed and incubated with 50 µL Biotinylated Complement C4 antibody per well for 1 hour. The microplate was washed again and incubated with 50 uL SP Conjugate per well for 30 minutes, followed by another plate wash, incubation with 50 µL Chromogen Substrate per well, and a final addition of 50 µL Stop solution. Absorbance was read on a Perkin Elmer Envision microplate reader at 450 nm.

**Fixed immunofluorescence sectioning, staining and imaging COs**

COs were prepped for harvesting, fixing, sectioning, and staining as described in protocol in Methods section. Primary antibodies used were rabbit EMX1 (1:100, Millipore-Sigma, HPA006421), goat SOX2 (1:500, R&D Systems, AF2018), rat CTIP2 (1:500, Abacm, ab18465) and mouse TBR1 (1:500, Protein Tech, 66564-1-Ig). Secondary antibodies and stains included donkey anti-rabbit AF647 (1:1000, Life-Technologies, A31573), donkey anti-goat CF750 antibody (1:1000, Millipore-Sigma, SAB4600444), donkey anti-rat CF750 (1:1000, Biotium, 20857), donkey anti-mouse AF488 (1:1000, Life Technologies, A21202), and DAPI (1:1000, Biovision, B1098-5).

**Supplemental Figures**

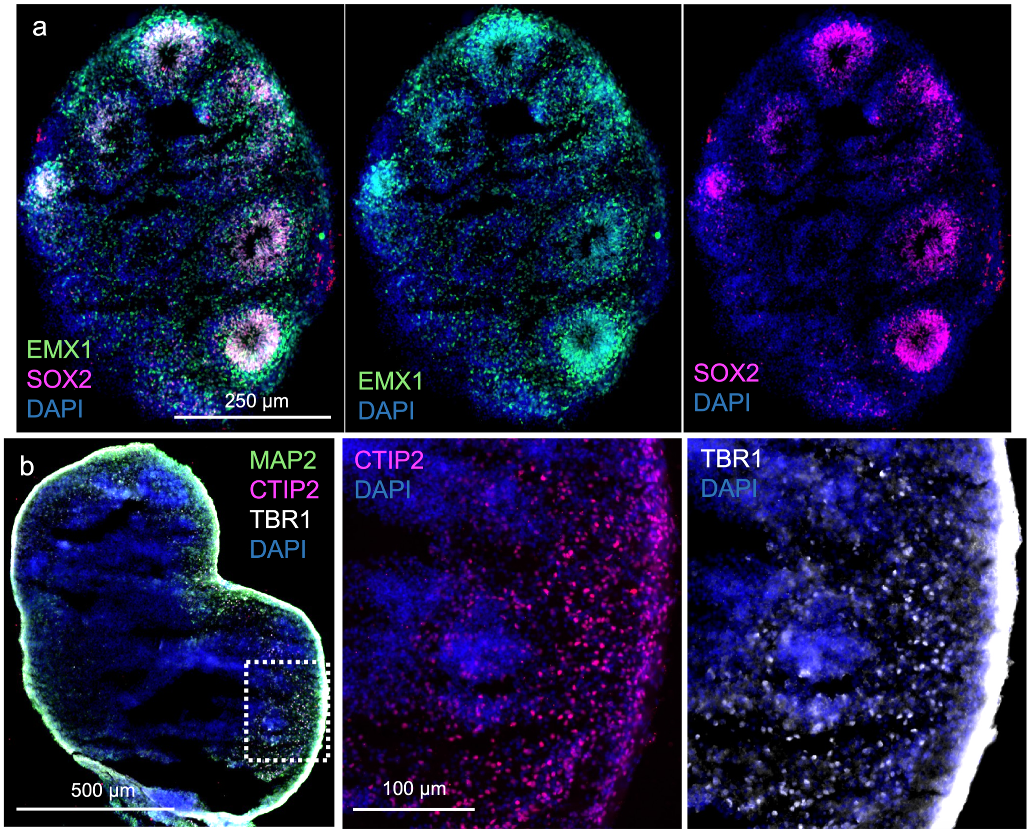


**Supplemental Figure 1.** (**a**) Day 60 CO section labeled with EMX1, SOX2, and DAPI. (**b**) Day 92 CO section labeled with MAP2, CTIP2, TBR1, and DAPI. White box indicates enlarged images from the CO slice.

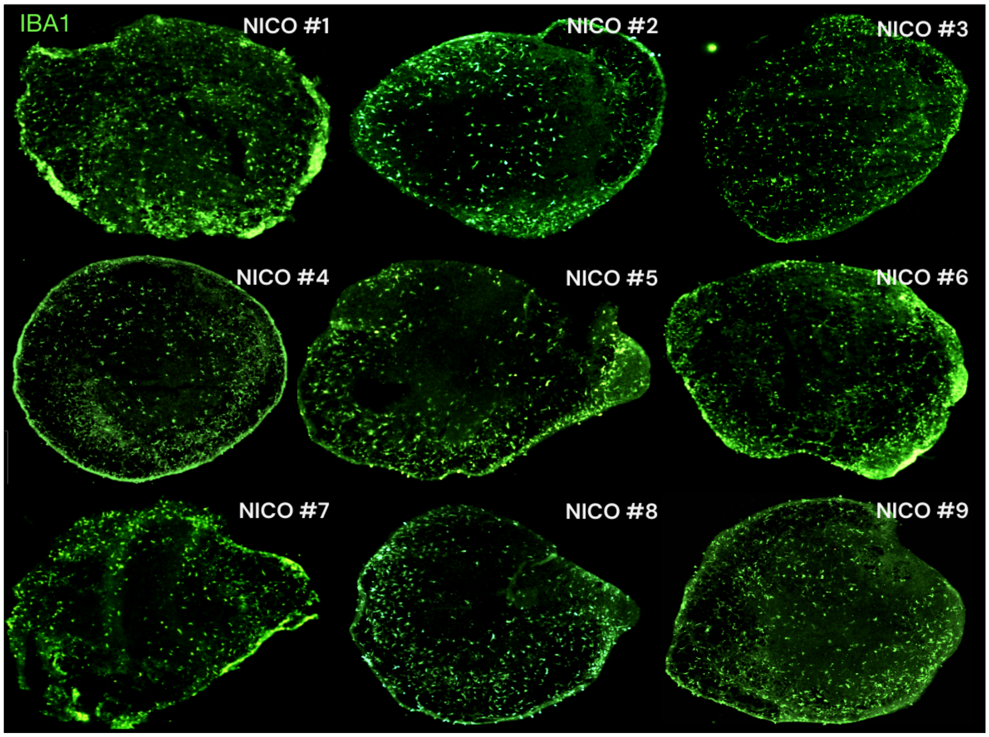


**Supplemental Figure 2.** NICO tissue sections demonstrating reproducibility of microglia integration across independent day 90 NICOs. NICO sections labeled with microglia marker IBA1.

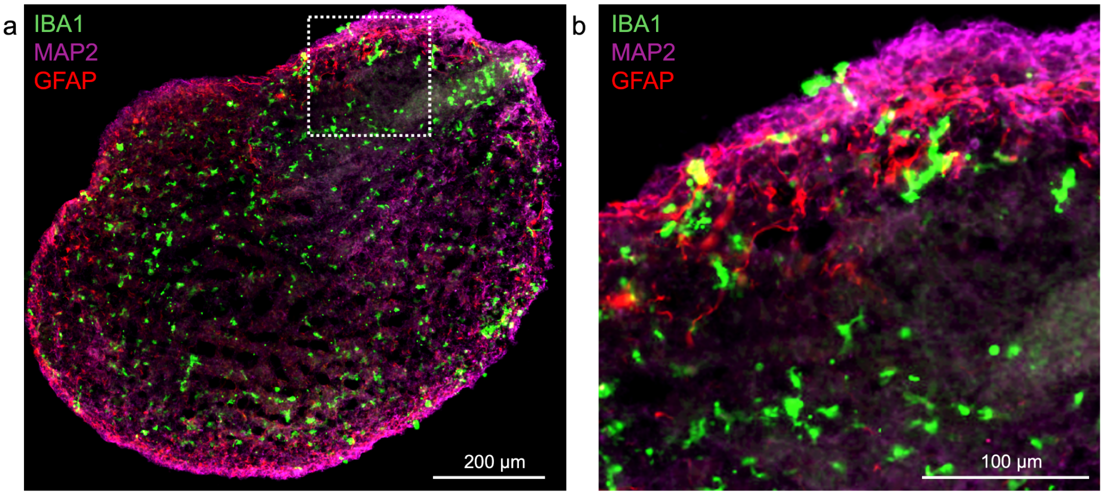


**Supplemental Figure 3.** NICO tissue sections demonstrating prolonged incubation of microglia 60 days post microglia integration. IBA1 labeled microglia are ramified and well distributed through the organoid body. NICOs were 131 days old at the time of harvest.

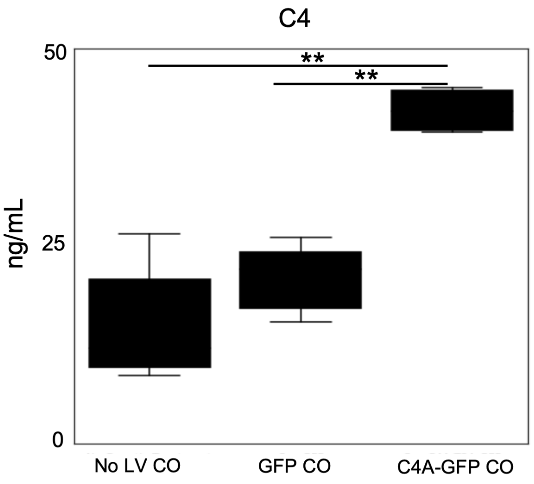


**Supplemental Figure 4.** Levels of secreted C4 in media from NICOs overexpressing C4A-GFP or GFP, or with No LV measured with C4 ELISA ($n$ = 3 independent media samples per parameter, box plot = upper and lower quartiles, with median center line, error bars = max and min of data set, statistical significance determined by one-way ANOVA with Tukey post-hoc tests, $^{**}p < 0.01$; corrected for multiple comparisons).

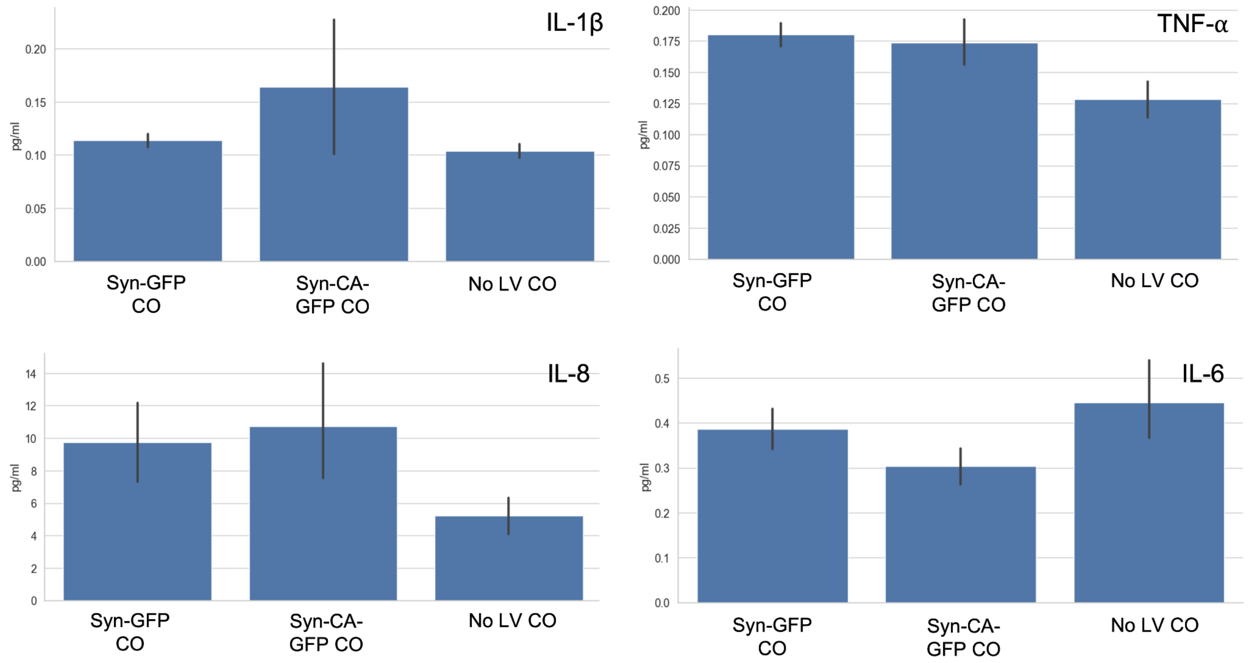


**Supplemental Figure 5.** Levels of secreted IL-1β, TNF-α, IL-6, and IL-8 cytokines in media from COs (without microglia) expressing GFP or C4A-GFP, or with No lentivirus (No LV), ($n$ = 8-10 independent samples per condition, bar = sample mean, error bar = SEM, statistical significance determined by one-way ANOVA with Tukey post-hoc tests, no significant difference in data between groups).

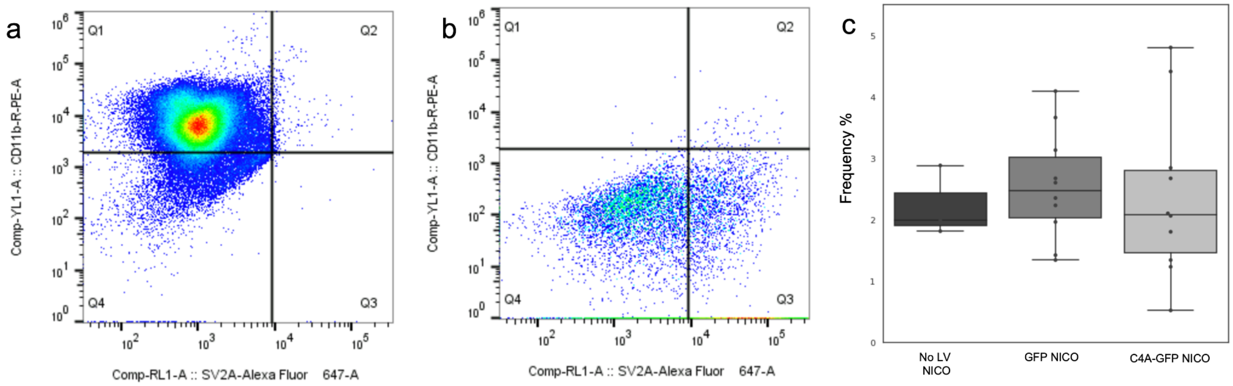


**Supplemental Figure 6.** Control groups for setting flow cytometry gates for CD11b-SV2A co-staining. (**a**) Microglia cells only without CO cells. (**b**) CO cells only without microglia cells. (**c**) Flow cytometry measured frequency (%) of CD11b-SV2A co-labeled cells from Q2 gate of No LV, GFP, and C4A-GFP NICOs (*n* = 4-12 separate organoids per parameter, box plot = upper and lower quartiles, with median center line, error bars = max and min of data set, statistical significance determined by one-way ANOVA with Tukey post-hoc tests, no significant difference in data between groups).

**Supplemental Table 1.** RT-qPCR primer sequences

| Primer | Forward seq | Reverse seq |
|---|---|---|
| GAPDH | TTGAGGTCAATGAAGGGGTC | GAAGGTGAAGGTCGGAGTCA |
| GFP | ACGTAAACGGCCACAAGTTC | AAGTCGTGCTGCTTCATGFG |
| C4A | CCCAATATGATCCCTGATGG | CCACTGCTCTGTCTTGTCCA |
| B2M | TGCTGTCTCCATGTTTGATGTATCT | TCTCTGCTCCCCACCTCTAAGT |
| CST7 | CGTCTGGATGACTGTGACTTCC | GGTCAGTGACAACGGAGAACAG |
| ITGAX | GATGCTCAGAGATACTTCACGGC | CCACACCATCACTTCTGCGTTC |
| LIPA | GTGGGTCATTCTCAAGGCACCA | CCATAGGGCTAGTACAGAAGGC |
| CX3CR1 | CACAAAGGAGCAGGCATGGAAG | CAGGTTCTCTGTAGACACAAGGC |
| LPL | CTGCTGGCATTGCAGGAAGTCT | CATCAGGAGAAAGACGACTCGG |
| CD9 | TCGCCATTGAAATAGCTGCGGC | CGCATAGTGGATGGCTTTCAGC |
| HLA | GCTGGAAACAGTTCCTCGGAGT | GACTCCACTCAGCATCTTGCTC |
| MERTK | CAGGAAGATGGGACCTCTCTGA | GGCTGAAGTCTTTCATGCACGC |
| SPP1 | CGAGGTGATAGTGTGGTTTATGG | GCACCATTCAACTCCTCGCTTTC |
| P2RY12 | TGCCAAACTGGGAACAGGACCA | TGGTGGTCTTCTGGTAGCGATC |
| APOE | CAACTCCTTCATGGTCTCGTCC | GGGTCGCTTTTGGGATTACCTG |
